# Genome-wide analysis exploring mechanisms used by *Shigella sonnei* to survive long-term nutrient starvation

**DOI:** 10.1101/2025.11.20.689435

**Authors:** Xosé M. Matanza, P. Brian Leung, Vincenzo Torraca, Jayne Watson, Matthew J. Dorman, Nicholas R. Thompson, Abigail Clements

## Abstract

*Shigella* is a major cause of severe diarrhoea with *S. flexneri* and *S. sonnei* accounting for over 90% of infections. As economies grow, *S. sonnei* replaces *S. flexneri* as the dominant cause of shigellosis, however the basis of this epidemiological shift remains unclear. Here we investigated whether *S. sonnei* is better equipped to survive nutrient starvation, a crucial condition for persistence both outside the host and within the colonic lumen. *S. sonnei* exhibited greater survival under long-term nutrient starvation (LTNS) than *S. flexneri*, rapidly activating survival mechanisms. We interrogated the genome of *S. sonnei* using Transposon Directed Insertion-site Sequencing (TraDIS) revealing that metabolic pathways (ATP, nucleotide, and amino acid synthesis), and envelope homeostasis complexes (e.g., Tol-Pal, Bam) are conditionally essential for LTNS. TraDIS findings were validated by non-competitive and competitive survival of wild-type and deletion mutant strains. We compared the homology of conditionally essential genes between *S. sonnei* and *S. flexneri* to identify putative genetic markers of differential interspecies LTNS survival. Analysis of *ldcA* (a peptidoglycan carboxypeptidase) and *rseA* (the anti-sigma factor regulator) indicated a major role in sustaining survival in LTNS in *S. sonnei*; however, allele-swap with *S. flexneri* alleles restored wild-type survival in *S. sonnei* suggesting that monogenic changes may not explain the divergent survival of these two species. Together, these data define the molecular adaptations of starvation resistance in *S. sonnei* and provide insights into its epidemiological dominance in high-income countries.

**Author summary:** Understanding why *S. sonnei* has a higher prevalence over *S. flexneri* as a country undergoes economic growth is one of the most important challenges in *Shigella* research. The investigation of their biological and genetic differences is key to tackle the impact of the disease. We discovered that *S. sonnei* resists nutrient deprivation better than *S. flexneri,* suggesting a better adaptation to an extracellular lifestyle and a greater preservation of metabolic capabilities. Using a genome-wide transposon sequencing approach we uncovered the key pathways behind the survival of *S. sonnei* facing nutrient starvation, which include ATP, nucleotide, and amino acid synthesis as well as maintenance of cell envelope integrity. Comparative analysis between *S. sonnei* and *S. flexneri* did not identify a single gene responsible for the differing survival and we suggest that the differing survival may stem from the coordination of multigenic differences. Our data provides a genome-wide basis for understanding how *S. sonnei* adapted to nutrient-deprived settings, which may be advantageous in the gut lumen and also in hostile environments, potentially contributing to its dominance in high-income countries.

## Introduction

*Shigella* spp. are Gram-negative bacteria of the order Enterobacterales that cause shigellosis or bacillary dysentery. Shigellosis is the second deadliest diarrhoeal disease, with 30% of fatalities occurring in children below five years of age (1). *Shigella* is highly contagious as only 10-100 cells suffice to cause disease (2), and spreads mainly via the faecal-oral route person-to-person (3–6) or by ingesting contaminated water or food (7–10). Fluoroquinolone-resistant *Shigella* is now considered by the World Health Organization (WHO) as a high-priority pathogen in need of urgent control measures (11). Despite research efforts, no licensed vaccine against shigellosis is currently available (12), although many are in development.

The genus *Shigella* includes four serogroups or species: *S. flexneri*, *S. sonnei*, *S. boydii* and *S. dysenteriae*. *S. flexneri* and *S. sonnei* cause more than 90% of episodes (13). Although *S. sonnei* infections are typically described as milder (14,15), comparative studies show that infections caused by both lineages manifest with similar severity, especially when infecting similar populations (16–18). Interestingly, the geographical distribution of *S. sonnei* and *S. flexneri* varies according to economic development: *S. flexneri* predominates in low-income countries (LIC) and *S. sonnei* in high-income countries (HIC) and middle-income countries (MIC). Importantly, global industrialisation and improved sanitation have been accompanied by a rise in *S. sonnei* infections (19–23). Epidemiological data suggest that hygiene measures that reduce *S. flexneri* infections in HIC are not as efficient at reducing *S. sonnei* (19). Understanding the underlying causes behind this phenomenon is therefore crucial. A prominent theory is related to natural immunisation: *S. sonnei* has a unique O-antigen that is identical to one serotype (O17) of the environmental bacterium *Plesiomonas shigelloides,* a bacterium that is widely distributed in aquatic environments (24,25). Exposure to the latter via contact with poorly sanitised waters is common in LICs and this could provide cross-protection against *S. sonnei* (26). Alternatively, ecological or physiological traits, including differences in interactions with the host or the microbiota, may explain the differing distributions.

*Shigella* is typically described as an intracellular pathogen. However we have previously shown that the O-antigen of *S. sonnei* reduces Type 3 Secretion System (T3SS) effectiveness resulting in less macrophage pyroptosis and reduced access to the cytoplasm of intestinal epithelial cells compared to *S. flexneri* (27). Additionally, unlike *S. flexneri*, *S. sonnei* encodes colicins, usually on the plasmid spB, that kill commensal members of the microbiota (28). In a zebrafish infection model, *S. sonnei* causes persistent infections with a high bacterial burden, while only certain *S. flexneri* serotypes persist but at lower bacterial burdens than seen for *S. sonnei* (29). These observations, coupled with its high level of antimicrobial resistance may give *S. sonnei* a competitive advantage in M/HIC (30).

Whilst most progress in bacteriology has focused on the study of organisms in nutrient abundance, bacteria are often exposed to periods of nutrient scarcity (31). If bacteria cannot acquire nutrients such as nucleobases, amino acids, and vitamins exogenously from the environment they must produce these nutrients through biosynthetic capabilities. How microbes modulate their physiology when living in starvation is a growing subject of research (32–34). *S. sonnei* can kill the competing microbiota and exhibits poor invasion of host cells. Therefore, we hypothesised that *S. sonnei* might also have evolved strategies to survive as an extracellular pathogen including an enhanced capacity to survive nutrient starvation. Here we confirm here that *S. sonnei* endures long-term nutrient starvation (LTNS) more effectively than *S. flexneri*, potentially providing *S. sonnei* with a survival advantage in highly sanitised environments (typical of wealthier countries) and in the colonic lumen (rather than intracellularly within the enterocyte cytosol). To comprehensively describe the molecular mechanisms used by *S. sonnei* to survive LTNS, we used a genome-wide approach using transposon-directed insertion site sequencing (TraDIS). TraDIS can be used to assess the genetic requirements for growth or survival under selected conditions such as nutrient scarcity (35–37). The pathways identified in our study as conditionally essential for survival in LTNS include nutrient and energy utilisation pathways and envelope homeostasis. We also used this data to attempt to identify why *S. sonnei* survived LTNS better than *S. flexneri*. However our results suggest that monogenic differences may not explain the distinct LTNS phenotype of these two *Shigella* species.

## Results

### S. sonnei is more resistant to nutrient starvation than S. flexneri

To assess if *S. sonnei* survives LTNS better than *S. flexneri,* we initially analysed the survival of two strains belonging to each of the two species (M90T and 2457T for *S. flexneri*, 381 and 53G for *S. sonnei*) in M9 media with no carbon source. As can be seen in Fig 1A, both *S. sonnei* strains maintain CFU numbers above 10^5^ CFU/ml over the long time-course. Conversely, *S. flexneri* strains decline abruptly from day 10 to a final concentration of 10^3^ CFU/ml at day 18. This finding led us to investigate how and when *S. sonnei* adapted to nutrient starvation. We supplemented *S. sonnei* M9 cultures with chloramphenicol, an antibiotic that targets ribosomal function to impede protein production. Chloramphenicol (Cm) was added at times 0, 2 and 24 h after exposure to M9 media with no carbon source. The number of viable cells was then monitored throughout the course of the experiment (Fig 1B). Only when Cm was added at 0 h did we observed a reduction in the number of viable cells. Interestingly the reduction did not occur until after day 7. When Cm was added at 2 h or 24 h, *S. sonnei* survival remained unaffected, suggesting that the changes necessary for survival are rapidly activated within the first 2 hours of starvation. Collectively, these results support our hypothesis and demonstrate that *S. sonnei* can rapidly adapt to nutrient deprivation and survive better than *S. flexneri*.

**Fig 1.**
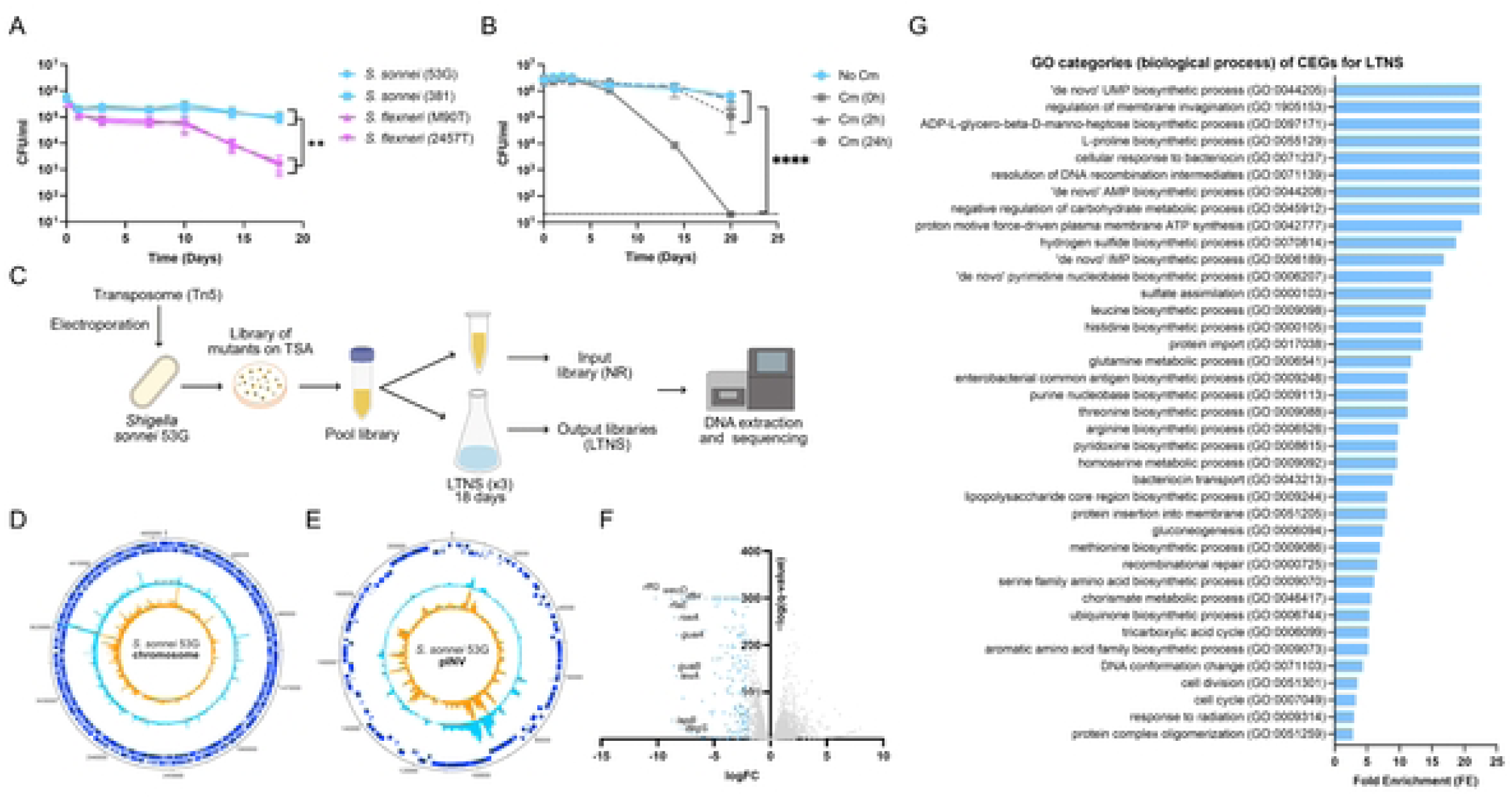
*Shigella sonnei* resists long-term nutrient starvation (LTNS) better than *Shigella flexneri*. TraDIS was used to identify the molecular mechanisms of *S. sonnei* to survive LTNS. A) Survival curves of *Shigella* strains including *S. sonnei* (53G and 381) and *S. flexneri* (M90T and 2457T) inoculated at an initial concentration of 10^6^ CFU/ml in minimal medium and monitored over 18 days. B) Survival curves of *S. sonnei* 381 in the presence of chloramphenicol (Cm) added at 0-, 2-, and 24-hours post-inoculation. Survival was compared to that in the absence of Cm and monitored over 20 days. For A and B, data show mean values ± SEM of n = 3 biological replicates. Log_10_-transformed data was analysed by 2-way ANOVA with the main column effect compared by Tukey’s multiple comparison. ** = p ≤ 0.01, **** = p ≤ 0.0001. C) Transposon-directed insertion site sequencing (TRADIS) was employed to decipher the genetic mechanisms involved in the survival of *S. sonnei* after long-term nutrient starvation (LTNS). A schematic representation of the TRADIS workflow is shown. A pool mutant library was generated in nutrient-rich (NR) conditions (input library) and then subjected to long-term nutrient starvation (LTNS) in triplicate (output libraries). The genomic DNA of both input and output libraries was extracted, sequenced and compared. D,E) Map of transposon insertion sites and frequency in input libraries (NR, orange) and output libraries (LTNS, blue) in *S. sonnei* 53G chromosome (D) and the LVP pINV (E). F) Volcano plot of LTNS TraDIS results showing fitness scores for genes in *S. sonnei* 53G chromosome showing their logFC vs adjusted p-value represented as -log(q-value). Conditionally essential genes (log FC >-2) are highlighted in blue. The 10 genes with the lowest logFC are indicated. G) Significant Gene Ontology categories of conditionally essential genes identified by TraDIS, using the PANTHER (Protein ANalysis THrough Evolutionary Relationships) Classification System. Only categories with a p-value < 0.05 are displayed.

### Transposon Directed Insertion-Site Sequencing (TraDIS) reveals LTNS survival mechanisms of S. sonnei

After confirming that *S. sonnei* is more resistant against nutrient depletion than *S. flexneri* we were intrigued to unveil the molecular mechanisms responsible. For this purpose, we designed a Transposon Directed Insertion-site Sequencing (TraDIS) approach to compare transposon insertion frequencies across the genome of *S. sonnei* grown in nutrient rich (NR) conditions (input library) or subjected to LTNS (output library) (Fig 1C). Firstly, we used transposon Tn5 to generate a comprehensive mutant library in the representative laboratory strain *S. sonnei* 53G grown in NR conditions. The library consisted of 136,857 independent insertions on the chromosome and 15,430 independent insertions on the LVP. This NR input library was subjected to LTNS and input/output libraries were sequenced and insertions mapped to the chromosome (Fig 1D) and the large virulence plasmid pINV (Fig 1E). Genes were classified according to the frequency of insertions per condition. 418 genes were determined to be essential; i.e, there were no insertions in input or output libraries (Supplementary Table 1). All the predicted essential genes were chromosomally encoded, with the exception of 2 short predicted pseudogenes (75 and 84 bp, respectively) located on pINV. Since pINV is known to be non-essential, these two hits are likely to represent false positives and may not have been covered in the transposon library because of their small size.

To uncover mechanistic approaches that go beyond universal essentiality, we focused our study on conditionally essential genes (CEGs) for LTNS, i.e., those with fewer insertions in the output than the input libraries (cut-off = LogFC <-2, p-value < 0.05) (Fig 1F). From the chromosome and the large virulence plasmid pINV (Fig 2B), bioinformatic analysis identified 218 CEGs. No conditionally essential or essential genes were identified on the remaining plasmids. The 50 genes with the most significant Log FC, are listed in Table 1. All CEGs genes can be found in Supplementary Table 2.

**Fig 2.**
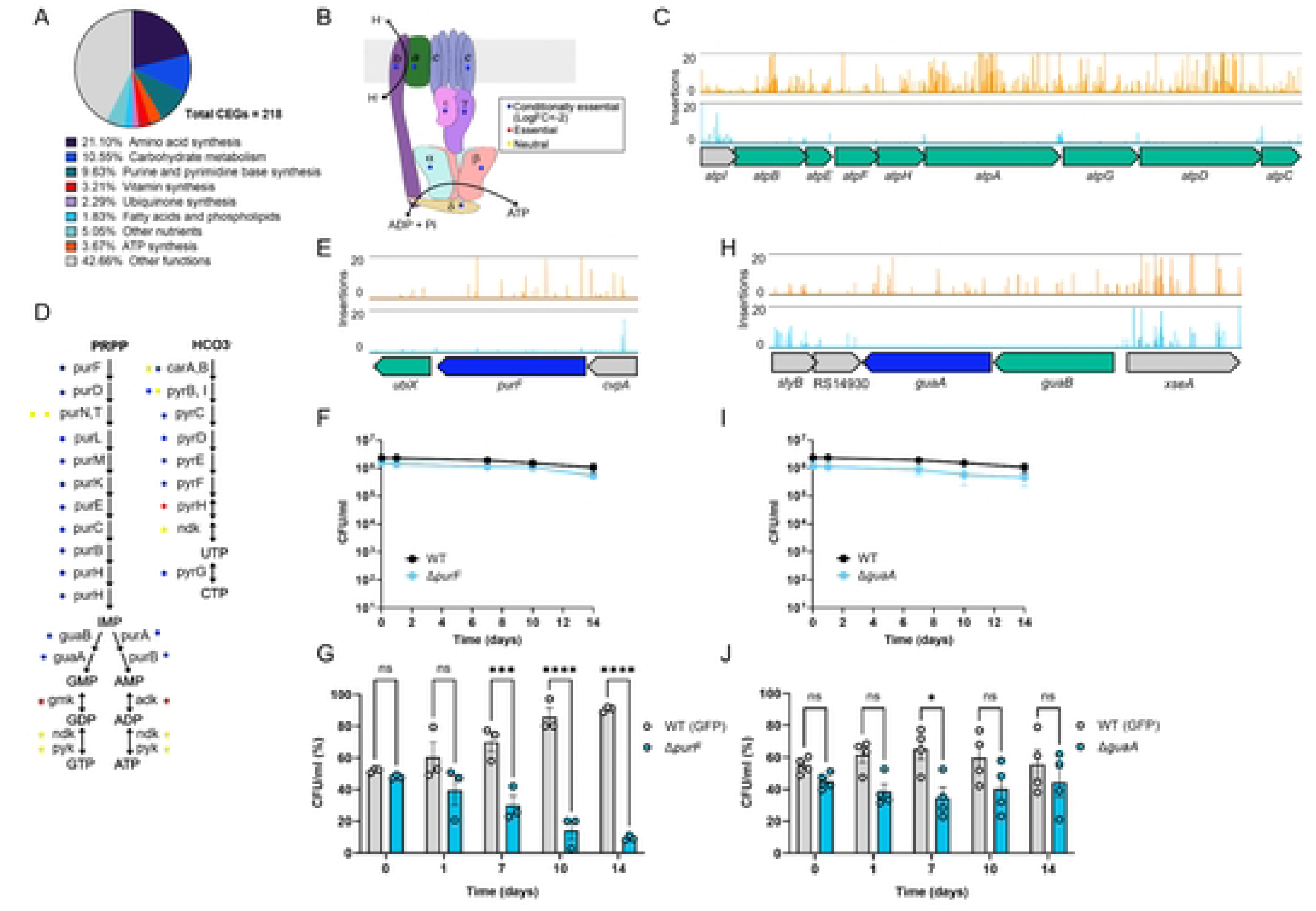
Transposon Directed Insertion site Sequencing (TraDIS) highlighted metabolic pathways including ATP and nucleotide synthesis pathways as conditionally essential for the survival of *S. sonnei* after long-term nutrient starvation (LTNS). A) Metabolic genes account for more than half of conditionally essential genes (CEGs) found in TRADIS analysis. B) ATP synthase subunits. Colour dots indicate their identification by TRADIS as conditionally essential (blue), essential (red) or neutral (yellow). C) Transposon insertions in ATP synthesis locus. Read counts are limited to 20. CEGs are coloured in green. D) Schematic of purine and pyrimidine synthesis pathways. Colour dots indicate gene identification by TRADIS as conditionally essential (blue), essential (red) or neutral (yellow). E) TRADIS insertions in *purF.* Read counts are limited to 20. F) Non-competitive LTNS survival of WT SS381 and SS381Δ*purF.* G) Competition assays between WT SS381 and SS381Δ*purF.* H) TRADIS insertions in *guaA.* Read counts are limited to 20. *purF* is coloured in blue and other CEGs are coloured in green. I) Non-competitive LTNS survival of WT SS381 and SS381Δ*guaA.* J) Competition assays between WT SS381 and SS381Δ*guaA.* For F and I, data show mean values ± SEM of n = 3 biological replicates. Log_10_-transformed data was analysed by 2-way ANOVA with main column effect compared by Tukey’s multiple comparison. For G and J data show mean values ± SEM at least 3 biological replicates. A two-way ANOVA with Šídák’s multiple comparisons was used to compare mean values at different time points. ns = non-significant (p > 0.05), * = p ≤ 0.05, *** = p ≤ 0.001, **** = p ≤ 0.0001.

**Table 1.** Top 50 genes, according to their LogFC, identified by TraDIS as conditionally essential for the survival of *S. sonnei* 53G under LTNS.

| Gene | Function | LogFC | p-value |
| --- | --- | --- | --- |
| *rffG* | dTDP-glucose 4,6-dehydratase | -10.20 | 0 |
| *wecD* | dTDP-4-amino-4,6-dideoxy-D-galactose acyltransferase | -8.82 | 0 |
| *YciM/LapB* | LPS assembly protein B | -8.67 | 3.82E-41 |
| *guaB* | IMP dehydrogenase | -8.55 | 2.29E-157 |
| *rseA* | anti-sigma-E factor RseA | -8.47 | 3.55E-261 |
| *lexA* | LexA repressor | -8.34 | 3.11E-136 |
| *guaA* | GMP synthase (glutamine-hydrolyzing) | -8.26 | 1.24E-224 |
| *degS* | outer membrane-stress sensor serine endopeptidase DegS | -7.95 | 1.00E-24 |
| *rfaE* | bifunctional heptose 7-phosphate kinase/heptose 1-phosphate adenylyltransferase | -7.94 | 0 |
| *rffH* | glucose-1-phosphate thymidyltransferase | -7.77 | 0 |
| *atpE* | ATP synthase subunit C | -7.75 | 4.43E-89 |
| *waaC* | lipopolysaccharide heptosyltransferase RfaC | -7.28 | 0 |
| *lepB* | S26 family signal peptidase | -7.14 | 3.25E-15 |
| *wecE* | dTDP-4-amino-4,6-dideoxygalactose transaminase | -7.14 | 0 |
| *waaF* | ADP-heptose--LPS heptosyltransferase 2 | -7.03 | 0 |
| *glmS* | glutamine--fructose-6-phosphate transaminase (isomerizing) | -6.92 | 7.03E-13 |
| *rfaD* | ADP-L-glycero-D-mannoheptose-6-epimerase | -6.72 | 0 |
| *atpD* | ATP synthase subunit beta | -6.59 | 0 |
| *atpA* | ATP synthase subunit alpha | -6.57 | 0 |
| *rpiA* | ribose-5-phosphate isomerase | -6.46 | 4.39E-09 |
| *secE* | protein translocase subunit SecE | -6.08 | 4.44E-07 |
| *atpF* | ATP synthase subunit B | -6.02 | 2.52E-83 |
| *ribA* | GTP cyclohydrolase II | -5.93 | 3.15E-06 |
| *wzyE* | O-antigen assembly polymerase | -5.93 | 3.15E-06 |
| *galU* | UTP--glucose-1-phosphate uridylyltransferase | -5.80 | 5.47E-233 |
| *pyrG* | CTP synthetase | -5.77 | 2.19E-05 |
| *surA* | chaperone SurA | -5.72 | 0 |
| *higA* | transcriptional regulator | -5.63 | 5.82E-20 |
| *tpiA* | triose-phosphate isomerase | -5.29 | 1.14E-59 |
| *gmhA* | phosphoheptose isomerase | -5.16 | 2.51E-53 |
| *cysQ* | 3'(2'),5'-bisphosphate nucleotidase CysQ | -5.14 | 0 |
| *rpsF* | 30S ribosomal protein S6 | -5.10 | 2.50E-04 |
| *proC* | pyrroline-5-carboxylate reductase | -5.09 | 2.31E-172 |
| *prc* | carboxy terminal-processing peptidase | -5.03 | 7.82E-62 |
| *purA* | adenylosuccinate synthetase | -5.02 | 0 |
| *atpG* | ATP synthase subunit gamma | -4.97 | 0 |
| *atpC* | ATP synthase epsilon chain | -4.94 | 1.03E-97 |
| *atpG* | ATP synthase subunit delta | -4.91 | 1.59E-141 |
| *atpA* | ATP synthase subunit A | -4.88 | 0 |
| *purK* | 5-(carboxyamino)imidazole ribonucleotide synthase | -4.80 | 3.69E-285 |
| *folP* | dihydropteroate synthase | -4.79 | 1.99E-03 |
| *purH* | bifunctional<br>phosphoribosylaminoimidazolecarboxamide<br>formyltransferase/inosine monophosphate<br>cyclohydrolase | -4.74 | 0 |
| *purD* | phosphoribosylamine--glycine ligase | -4.69 | 1.39E-172 |
| *wecF* | TDP-N-acetylfucosamine:lipid II N-acetylfucosaminyltransferase | -4.68 | 0 |
| *carB* | carbamoyl phosphate synthase large subunit | -4.65 | 0 |
| *pyrB* | aspartate carbamoyltransferase catalytic subunit | -4.59 | 1.28E-293 |
| *acnB* | aconitate hydratase B | -4.55 | 0 |
| *recB* | exodeoxyribonuclease V subunit beta | -4.51 | 0 |
| *purM* | phosphoribosylformylglycinamide cyclo-ligase | -4.43 | 0 |
| *purC* | phosphoribosylaminoimidazolesuccinocarboxamide<br>synthase | -4.42 | 1.05E-132 |

### Analysis of conditionally essential genes and pathways

To uncover the main pathways that sustain LTNS survival in *S. sonnei*, we assigned the set of CEGs to GO categories (biological process) (Fig 1G and Supplementary Table 3). 39 significant GO categories (p-value < 0.05) were obtained. Fewer mutants were recovered in genes related to nutrient metabolism and energy acquisition. Pathways with a high Fold Enrichment (FE) include *’de novo’* UMP biosynthetic process (GO:0044205), L-proline biosynthetic process (GO:0055129), proton motive force-driven plasma membrane ATP synthesis (GO:0042777), leucine biosynthetic process (GO:0009098), and histidine biosynthetic process (GO:0000105).

Our results reveal that among the CEGs *S. sonnei* requires to survive LTNS, more than half (n=125/218) participate in the metabolism of major nutrients and energy: carbohydrates, amino acids, nucleotides, nitrogenous bases, vitamins and ATP (Fig 2A). These findings not only align with expectations and reinforce the validity of our experimental approach, but they also reveal the critical metabolic requirements for *S. sonnei* facing nutrient deprivation. Genes encoding enzymes that participate in the central metabolism of carbohydrates were found to be conditionally essential: these include key glycolysis enzymes such as *pgm* (phosphoglucomutase) and *pgi* (phosphoglucose isomerase) along with several enzymes of the tricarboxylic acid cycle, including *acnB* (aconitase B), *gltA* (citrate synthase), *sucA*, and *sucB* (components of the 2-oxoglutarate dehydrogenase complex). Glycolysis produces pyruvate that can be incorporated into the tricarboxylic acid cycle to produce energy and precursors for the synthesis of various biomolecules. Additionally, genes involved in the synthesis of glycogen from glucose (*glpD, glgA, glgC*) and gluconeogenesis genes (*tpiA, pgi, fbp*) were found among this set of genes, suggesting that the production of glycogen from non-carbohydrate sources is also vital under starvation conditions. The synthesis of at least 13 amino acids (methionine, threonine, leucine, isoleucine, valine, proline, tryptophan, histidine, cysteine, tyrosine, arginine, glutamine and serine) was found to be conditionally essential. Likewise, the majority of genes involved in the biosynthesis of nitrogenous bases, both purines and pyrimidines, together with the locus that encodes for the ATP synthase complex were found among this set. Genes encoding for nucleotide and amino acid synthesis functions were selected for individual testing to validate TraDIS findings.

### Metabolic pathways of purine and amino acid synthesis are crucial for the survival of S. sonnei in LTNS

We constructed deletion mutants of selected CEGs in *S. sonnei* strain SS381 and assessed the effect of these mutations by comparing survival in single-strain cultures but also in competition assays with the wild-type (WT) strain. We chose to use SS381, given that this strain demonstrates the same survival as the 53G strain during LTNS (Fig 1A), and is representative of a more epidemiologically-prevalent lineage of *S. sonnei* (i.e., Lineage 3). To discriminate between WT and mutant strains during competition assays, we introduced a gene encoding green fluorescent protein (GFP) downstream of *glmS* (glucosamine-fructose-6-phosphate aminotransferase) in SS381, a site of insertion previously identified as neutral (38). We ensured that this *gfp* insertion had no effect on SS381 survival during LTNS and could therefore be used as the WT (Supplementary Fig 1). For comparison, the growth of all strains used in this study in NR conditions can be found in Supplementary Fig 2.

Since most genes encoding for the ATP synthase complex were identified as conditionally essential (Fig 2B) and this complex has a central role in energy conservation, we initially targeted genes *atpA* and *atpE* for mutagenesis. Despite numerous attempts we were unable to obtain deletions of these genes, even though *atpA* and *atpE* were identified as conditionally essential in contrast to essential (Fig 2C). However, we were successful in the construction of a deletion mutant in *purF.* Many genes that participate in the synthesis of purines and pyrimidines were found as CEGs (Fig 2D). PurF (amidophosphoribosyltransferase) initiates the purine biosynthesis pathway by converting 5-phosphribosyl-a-1-pyrophosphate to 5-phosphoribosylamine. This pathway produces inosine monophosphate (IMP), the precursor of adenine and guanine nucleotides (Fig 2D and 2E). Δ*purF* survival was assessed in single-strain cultures and in our competition assay. In single-strain cultures the difference in survival between WT and Δ*purF* was not significant while in the competition assay Δ*purF* demonstrated significant survival defects compared to the WT strain from 7 days post starvation (Fig 2F and 2G). We also constructed a mutant in *guaA*, a guanosine monophosphate (GMP) synthetase which catalyses the last step of GMP synthesis (Fig 2D and 2H). In single-strain cultures Δ*guaA* did not show any impairment in comparison to the WT, while in the competition assay it showed impairment but was only significant at day 7 (Fig 2I and Fig 2J). This suggests that de-novo nucleotide synthesis is important for LTNS survival.

We then constructed deletion mutants in selected amino acid synthesis pathways: methionine (*metE*), and cysteine (*cysI*). These mutants were tested in our single-strain survival and competition assays to determine their conditional essentiality during LTNS. MetE, a cobalamin-independent homocysteine transmethylase, catalyses the final step of methionine biosynthesis (Fig 3A and 3B). Δ*metE* showed no impairment in the single-strain culture but was impaired in LTNS survival in comparison to the WT at early time points (day 1). At later timepoints however, the survival levels were comparable to those of the WT and may be due to MetH, the cobalamin-dependent homocysteine transmethylase compensating for the MetE deletion (Fig 3C and 3D). CysI and CysJ are subunits of the sulfate reductase that produces sulfate necessary for cysteine synthesis (Fig 3E and 3F). A deletion in *cysI* again showed no impairment in the single-strain culture but did have significantly reduced survival compared to WT from day 7 onwards during the competition assay (Fig 3G and 3H).

**Fig 3.**
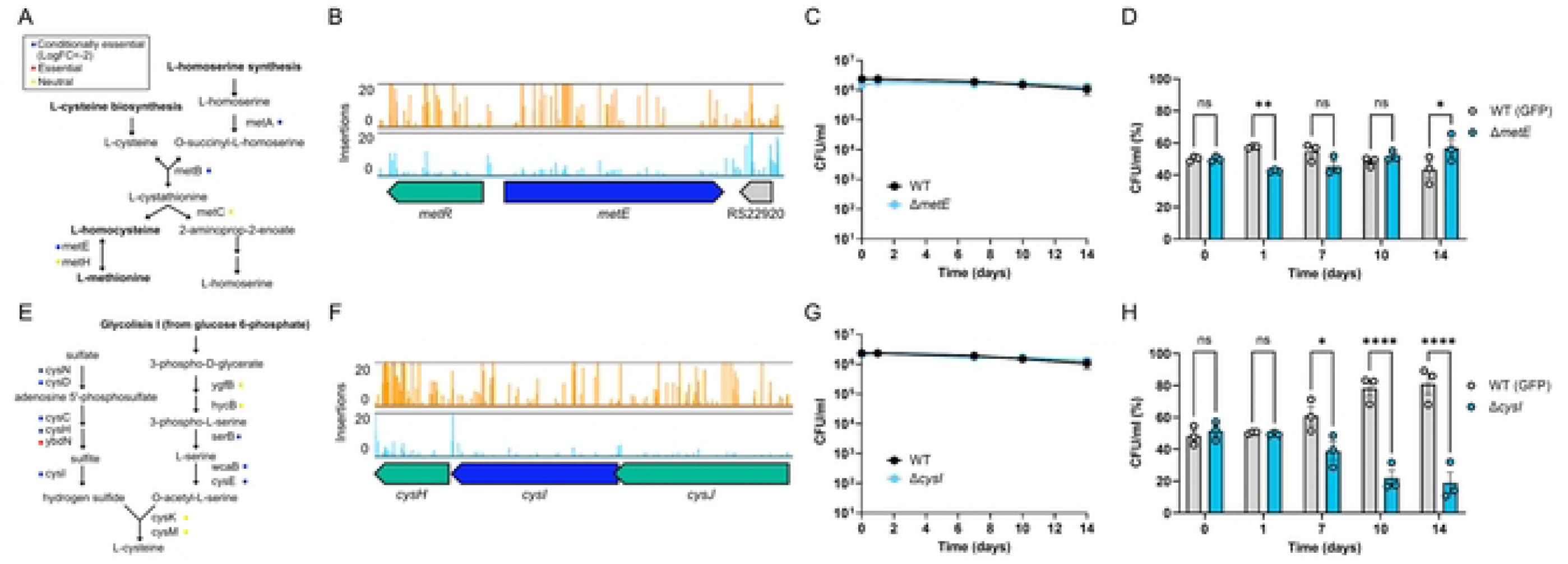
Multiple amino acid synthesis pathways are conditionally essential during long-term nutrient starvation (LTNS). A) Schematic of methionine synthesis pathway. Colour dots indicate gene identification by TRADIS as conditionally essential (blue), essential (red) or neutral (yellow). B) Transposon insertions in *metE*. Read counts are limited to 20. *metE* is coloured in blue and other CEGs are coloured in green.C) Non-competitive LTNS survival of WT SS381 and SS381Δ*metE.* G) Competition assays between WT SS381 and SS381Δ*metE.* E) Cysteine synthesis pathway. Colour dots indicate the identification of genes by TRADIS as conditionally essential (blue), essential (red) or neutral (yellow). F) Transposon insertions in *cysI*. Read counts are limited to 20. *cysI* is coloured in blue and other CEGs are coloured in green. G) Non-competitive LTNS survival of WT SS381 and SS381Δ*cysI.* H) Competition assays between WT SS381 and SS381Δ*cysI.* For C and G, data show mean values ± SEM of n = 3 biological replicates. Log_10_-transformed data was analysed by 2-way ANOVA with main column effect compared by Tukey’s multiple comparison. For D and H, data show mean values ± SEM of n=3 biological replicates. A two-way ANOVA with Šídák’s multiple comparisons was used to compare mean values at different time points. ns = non-significant (p > 0.05), * = p ≤ 0.05, ** = p ≤ 0.01, **** = p ≤ 0.0001.

Together, these results validate our TRADIS findings highlighting nucleotide and amino acid synthesis pathways as important for *S. sonnei* to survive LTNS. These pathways are crucial for maintaining the nucleotide pools required for DNA and RNA synthesis and repair, and the amino acid pool for protein synthesis under nutrient-restricted conditions. Furthermore, we demonstrate the sensitivity of our competition assay in determining survival differences between strains.

### Cell-envelope homeostasis has a role in the survival of S. sonnei in LTNS

Most CEGs identified in this study belong to nutrient and metabolic pathways, however, categories related to envelope homeostasis were also highlighted (Fig 1G). Genes involved in the synthesis and regulation of molecules present in the cell envelope were also substantially represented in categories such as regulation of membrane invagination (GO:1905153), enterobacterial common antigen biosynthetic process (GO:0009246), lipopolysaccharide core region biosynthetic process (GO:0009244), and protein insertion into membrane (GO:0051205).

The five proteins encoding for the Tol-Pal transmembrane complex were conditionally essential for survival in LTNS. This system participates in membrane invagination, removes excessive PL from the OM, and ensures envelope stability by linking the OM, PG and IM (62–65). TolA extends across the periplasm, connecting the OM and IM components of the system (Fig 4A and 4B). We constructed a deletion mutant of *tolA* and performed a competition assay with WT. Δ*tolA* had reduced survival from day 1 of LTNS when in competition with WT, although this was only statistically significant at day 1 and day 10 (Fig 4C and 4D).

**Fig 4.**
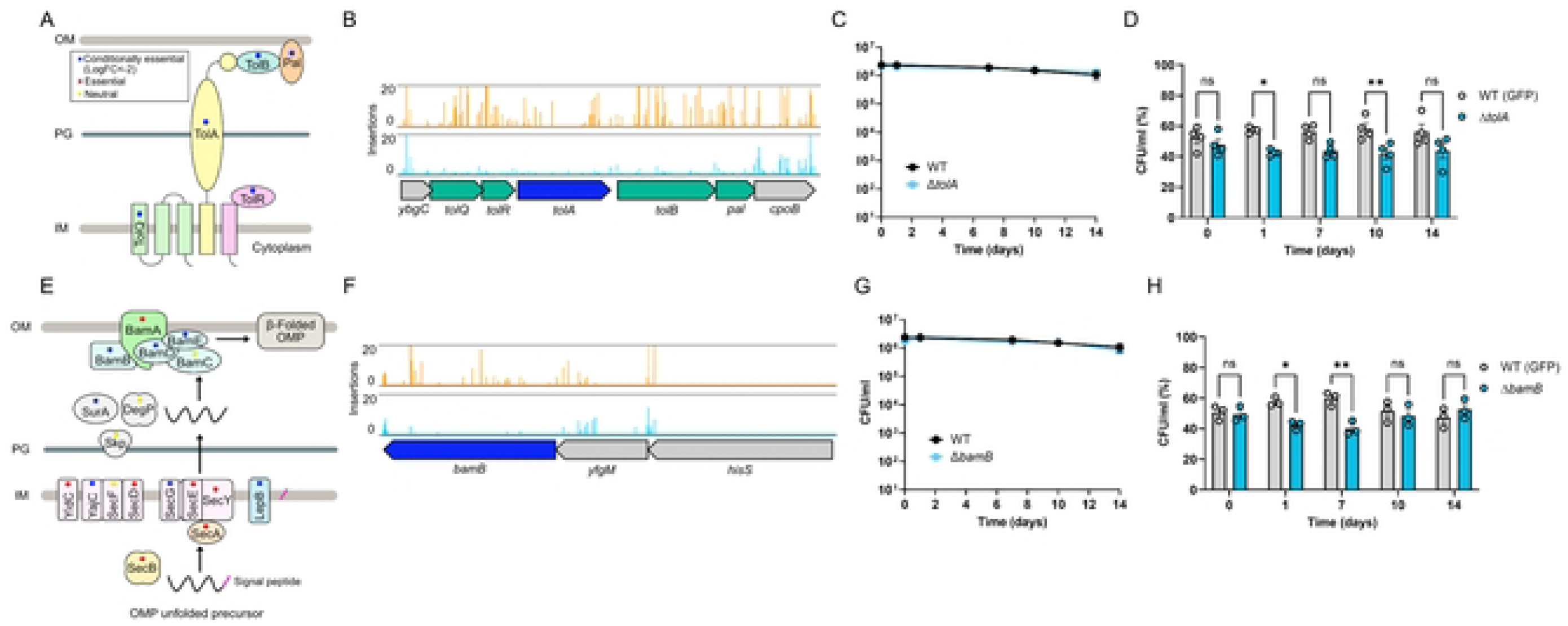
Maintaining cell envelope homeostasis contributes to long-term nutrient starvation (LTNS) survival. A) Schematic representation of the Tol-Pal system in the cell envelope. Colour dots indicate gene identification by TRADIS as conditionally essential (blue), essential (red) or neutral (yellow). B) Transposon insertions in genes encoding for the Tol-Pal complex. Read counts are limited to 20. *tolA* is coloured in blue and other CEGs are coloured in green. C) Non-competitive LTNS survival of WT SS381 and SS381Δ*tolA.* D) Competition assays between WT SS381 and SS381Δ*tolA.* E) Schematic representation of the Bam complex in the cell envelope. Colour dots indicate gene identification by TRADIS as conditionally essential (blue), essential (red) or neutral (yellow). F) Transposon insertions in *bamB*. Read counts are limited to 20. *bamB* is coloured in blue and other CEGs are coloured in green. G) Non-competitive LTNS survival of WT SS381 and SS381Δ*bamB.* H) Competition assays between WT SS381 and SS381Δ*bamB.* For C and G, data show mean values ± SEM of n = 3 biological replicates. Log_10_-transformed data was analysed by 2-way ANOVA with main column effect compared by Tukey’s multiple comparison. For D and H, data show mean values ± SEM of at least 3 biological replicates. A two-way ANOVA with Šídák’s multiple comparisons was used to compare mean values at different time points. ns = non-significant (p > 0.05), * = p ≤ 0.05, ** = p ≤ 0.01.

Outer membrane proteins (OMPs) are generated in the cytoplasm and transported to the OM (Fig 4E). Initially, unfolded OMPs cross the IM with the aid of the Sec system directed by their amino-terminal signal peptide. The Sec system is ubiquitous in bacteria and is formed by the SecYEG translocon, auxiliary proteins SecDF-YajC-YidC and chaperones SecAB (58). Following cleavage of the signal peptide by LepB, OMPs are transported to the OM by periplasmic chaperones (SurA, Skp, DegP, FkpA) (59). OMPs are then folded in their β-barrel conformation and inserted into the OM by the β-barrel assembly machinery (BAM) complex, made up of 5 proteins, BamA anchored in the OM and lipoproteins BamB, BamC, BamD and BamE (60). Our analysis found the majority of genes in these pathways to be either conditionally essential (SecG, YajC, LepB, SurA, BamB and BamD) or essential for viability (ie. no insertions found in any study condition, SecY, SecB, YidC and BamA). We selected *bamB* for deletion and characterisation under LTNS (Fig 3F). Δ*bamB* survived in single-strain cultures at a level equivalent to WT, while in the competition assay Δ*bamB* had an impaired phenotype at early timepoints, while at the later timepoints Δ*bamB* recovery was equivalent to WT (Fig 3G and 3H).

### Identifying genes responsible for the difference between S. sonnei and S. flexneri survival of LTNS

To investigate genetic candidates that could explain the difference in LTNS survival between *S. flexneri* and *S. sonnei,* we compared the protein sequence of each CEG identified in *S. sonnei* 53G with their counterpart in *S. flexneri* M90T and 2457T. Genes of interest were those absent or non-homologous in the *S. flexneri* genomes (Supplementary Table 4). We further ensured that the putative candidates were conserved in *S. sonnei* 381 (Fig 5A and 5B). This analysis identified one promising candidate that was absent from both *S. flexneri* genomes and 100% identical in the *S. sonnei* genomes: *yncJ* (Fig 5C). YncJ is a largely uncharacterised protein that contains a domain of unknown function (DUF2554) and appears to be restricted to *Enterobacteriaceae*. We constructed a *ΔyncJ* strain and analysed survival in our LTNS assays. However this mutant did not show a significantly decreased survival in LTNS compared to WT in either single-strain cultures or in competition assays (Fig 5D and 5E).

**Fig 5.**
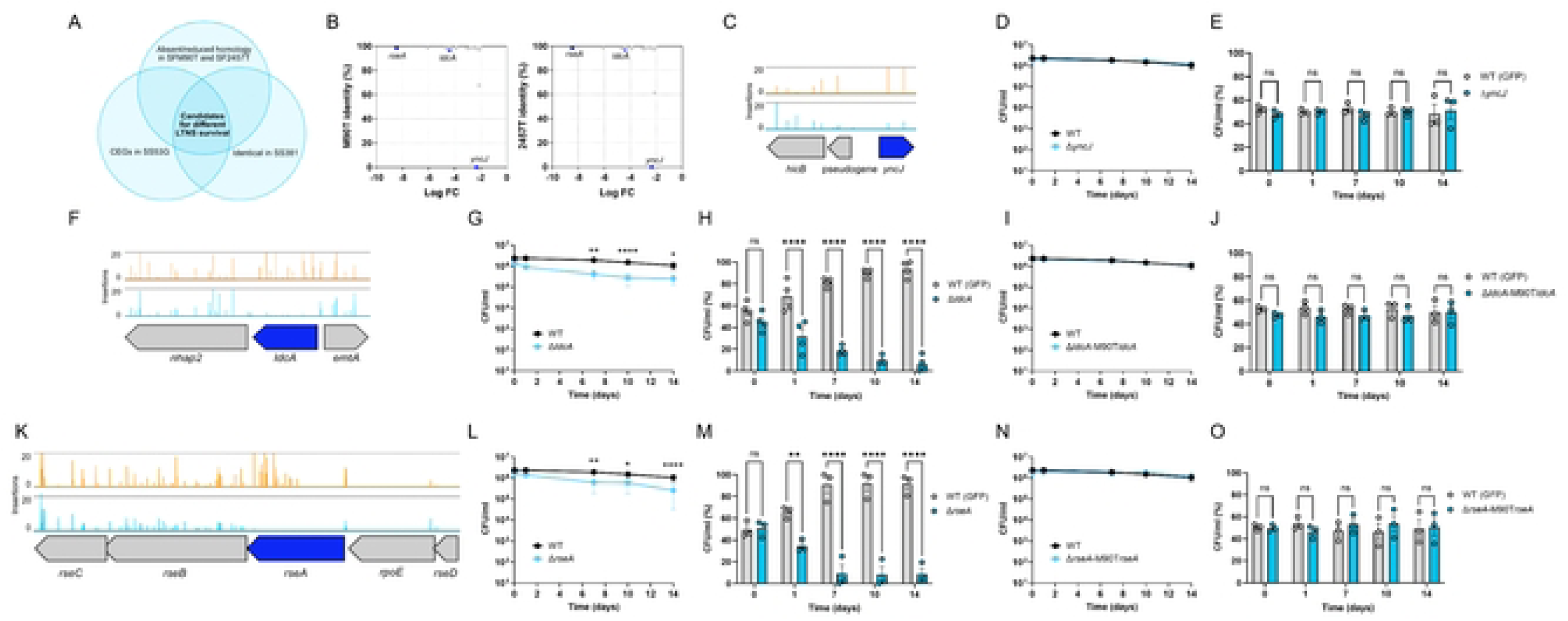
Identification of potential genetic markers that explain the different ability of *S. sonnei* and *S. flexneri* to resist long-term nutrient starvation (LTNS). A) Venn Diagram showing the chosen criteria for the identification of genetic candidates potentially explaining the different ability of *S. sonnei* and *S. flexneri* to resist LTNS: being conditionally essential in *S. sonnei* 53G, showing absent or reduced homology in *S. flexneri* M90T and *S. flexneri* 2457T, and being identical in *S. sonnei* 381. B) 24 Genetic candidates sharing <99% identity with both *S. flexneri* M90T and *S. flexneri* 2457T according to their log FC and amino acid identity. C) Transposon insertions in *yncJ*. Read counts are limited to 20. *yncJ* is coloured in blue and other CEGs are coloured in green. D) Non-competitive LTNS survival of WT SS381 and SS381Δ*yncJ.* E) Competition assays between WT SS381 and SS381Δ*yncJ.* F) Transposon insertions in *ldcA*. Read counts are limited to 20. *ldcA* is coloured in blue and other CEGs are coloured in green. G) Non-competitive LTNS survival of WT SS381 and SS381Δ*ldcA.* H) Competition assays between WT SS381 and SS381Δ*ldcA.* I) Non-competitive LTNS survival of WT SS381 and SS381Δ*ldcA-*M90T*ldcA.* J) Competition assays between WT SS381 and SS381Δ*ldcA-*M90T*ldcA.* K) Transposon insertions in *rseA*. Read counts are limited to 20. *rseA* is coloured in blue and other CEGs are coloured in green. L) Non-competitive LTNS survival of WT SS381 and SS381Δ*rseA.* M) Competition assays between WT SS381 and SS381Δ*rseA.* N) Non-competitive LTNS survival of WT SS381 and SS381Δ*rseA-*M90T*rseA.* O) Competition assays between WT SS381 and SS381Δ*rseA-*M90T*rseA.* For D, G, I, L, and N data show mean values ± SEM of n = 3 biological replicates. Log_10_-transformed data was analysed by 2-way ANOVA with main column effect compared by Tukey’s multiple comparison. For E, H, J, M, and O data show mean values ± SEM of at least 3 biological replicates. A two-way ANOVA with Šídák’s multiple comparisons was used to compare mean values at different time points. ns = non-significant (p > 0.05), * = p ≤ 0.05, ** = p ≤ 0.01, **** = p ≤ 0.0001.

We then focused on genes that were non-homologous between the *S. sonnei* and *S. flexneri* genomes. 109 CEGs met our criteria (100% amino acid identity in *S. sonnei* genomes, <100% amino acid identity in both *S. flexneri* genomes). In order to choose candidates that could be responsible for the difference between *S. sonnei* and *S. flexneri* we combined information on the homology between the species (looking for non-conservative aa changes) and the logFC from TraDIS. We identified 24 candidates that share <99% identity with both M90T and 2457T (Table 2, Fig 5B). We chose two candidates to focus on: *ldcA*, which had a high logFC and several non-conservative aa changes and *rseA* which had the highest logFC, and one non-conservative amino acid change.

**Table 2.** CEGs identified in *S. sonnei* 53G that show less than 99% identity with both *S. flexneri* M90T and 2457T.

| Gene | Log FC | M90T identity (%) | 2457T identity (%) |
| --- | --- | --- | --- |
| *yncJ* | -2.33 | 0 | 0 |
| *ybcO* | -2.13 | 67.71 | 61.22 |
| *ldcA* | -4.39 | 97.04 | 97.36 |
| *frsA* | -2.10 | 97.06 | 0 |
| *yejL* | -2.12 | 97.33 | 97.33 |
| *hisIE* | -2.90 | 98.03 | 97.54 |
| *Int* | -2.94 | 98.05 | 97.66 |
| *rfaP* | -2.40 | 98.11 | 98.11 |
| *rfaH* | -2.31 | 98.15 | 98.15 |
| *ruvC* | -3.07 | 98.27 | 98.27 |
| *pyrF* | -3.96 | 98.37 | 97.96 |
| *secE* | -6.08 | 98.43 | 98.43 |
| *ubiH* | -2.25 | 98.47 | 98.47 |
| *pyrE* | -4.40 | 98.59 | 98.59 |
| *purD* | -4.69 | 98.6 | 98.37 |
| *glpD* | -3.88 | 98.6 | 98.76 |
| *ilvY* | -2.83 | 98.65 | 98.65 |
| *rseA* | -8.47 | 98.71 | 98.61 |
| *hemC* | -3.84 | 98.72 | 98.72 |
| *cysN* | -2.18 | 98.74 | 98.95 |
| *pdxJ* | -2.35 | 98.77 | 98.77 |
| *moeB* | -2.05 | 98.8 | 98.8 |
| *cysJ* | -2.63 | 98.83 | 98.83 |
| *infB* | -2.04 | 98.88 | 98.99 |

LdcA (Fig 5F) is a carboxypeptidase that participates in PG recycling by releasing the terminal D-alanine residue from the cytoplasmic tetrapeptide allowing reuse of the tripeptide by the former (39). There are 9 amino acid differences between *S. sonnei* and *S. flexneri* LdcA, 3 of which are non-conservative (Supplementary Fig 3A). Δ*ldcA* had significantly reduced survival in LTNS compared to WT in both the single-strain culture from day 7 (Fig 5G) and competition assays from 1 day post starvation, indicating a requirement for survival in LTNS (Fig 5H). This finding infers that PG recycling is key to survive starvation, potentially due to a reduction in *de novo* PG synthesis. We then complemented Δ*ldcA* with the M90T *ldcA* and compared this to WT in non-competitive conditions and in a competition assay. Unexpectedly, the M90T *ldcA* fully complemented Δ*ldcA* with no significant difference at any timepoint between the WT and Δ*ldcA-*M90T*ldcA* in both experiments (Fig 5I and 5J). Therefore, differences between *S. flexneri* and *S. sonnei* alleles of *ldcA* alone do not explain the difference in LTNS survival between these species.

We then tested the importance of RseA (Fig 5K) in surviving LTNS. *rseA* ranks as the fifth most impaired gene in our screening with a logFC of -8.47. RseA is the anti-sigma factor of σ^E^, a key regulator of the extracytoplasmic stress response and one of the sensory pathways (together with Cpx, Rcs, and Psp) that respond to envelope stress (40). There are 3 amino acid differences between *S. sonnei* and *S. flexneri* RseA, 1 of which is non-conservative and within the NT σ^E^ binding region (Supplementary Fig 3B) (41). In single-strain cultures there was a significant difference in survival between WT and Δ*rseA* from day 7 post starvation (Fig 5L), while in the competition assay Δ*rseA* had significant survival defects compared to the WT strain from 1 day post starvation (Fig 5M). Complementation with M90T *rseA* gave similar results to those with *ldcA*, no significant differences were observed between the WT and the complemented strains (Fig 5N and 5O). These results reinforce the importance of cell envelope homeostasis for LTNS survival in *S. sonnei*. However, the cause for the difference in LTNS survival between *S. flexneri* and *S. sonnei* could not be attributed to a single genetic factor.

## Discussion

The epidemiological landscape of shigellosis is evolving, with *S. sonnei* progressively replacing *S. flexneri* in countries undergoing economic growth. Therefore, understanding the differences between these two species is crucial to mitigating the impact of the disease. Here we show that *S. sonnei* is more resistant to LTNS than *S. flexneri,* potentially contributing to the dominance of *S. sonnei* in HIC. Environmental transmission of *S. sonnei* is thought to be lower than that of *S. flexneri* (42), and while this seems plausible in LIC, evidence suggests an increase of potentially environmentally transmitted *S. sonnei* infections in HIC (43). Since improved water sanitation reduces faecal contamination and nutrient levels in water, the ability of *S. sonnei* to survive LTNS could provide an advantage compared to *S. flexneri* in highly sanitised aquatic environments typical of HIC.

*Shigella* spp. are well documented to have undergone genome degradation reducing the metabolic capabilities compared to the ancestral *E. coli*. Among *Shigella* spp., metabolic modelling studies indicate that *S. sonnei* has retained more metabolic capabilities than *S. flexneri* (44,45), which is consistent with our results. The enhanced LTNS resistance of *S. sonnei* may allow it to survive extended periods extracellularly within the colonic lumen or within the environment. Surviving nutrient limitation could compensate for the reduced intracellular invasion levels of *S. sonnei* (mediated by the O-Ag in the LPS and the capsule) (27) by reducing the dependence on cell invasion for the acquisition of nutrients. The commensal microbiota restricts pathogen colonisation through nutrient limitation (46) and the ability to survive LTNS, together with the expression of colicins by most *S. sonnei* clinical isolates (28), could be advantageous for *S. sonnei* to survive in the gut lumen. Surviving starvation can also result in persistent infections (47), therefore our findings may contribute to explaining the observation that *S. sonnei* causes persistent infections more efficiently than *S. flexneri* in the zebrafish infection model (29).

We applied a genome-wide TraDIS approach to uncover the genetic mechanisms important for LTNS survival in *S. sonnei*. We identified 218 genes as conditionally essential, and more than 50% of those participate in metabolic pathways such as the synthesis of ATP, nucleotides, and amino acids. Interestingly many of the pathways identified previously as important for Enterohaemorrhagic *E. coli* (EHEC) colonisation of the gastrointestinal tract were also identified in our study: purine, pyrimidine, methionine, threonine, leucine, isoleucine, valine, proline, tryptophan, histidine, tyrosine, arginine, glutamine and serine synthesis (48,49). These convergences highlight the significant overlap between pathways required for surviving nutrient starvation and those required for surviving gastrointestinal colonisation, supporting again our hypothesis that *S. sonnei* may retain metabolic pathways that support survival in the gastrointestinal tract.

Our data also highlighted the importance of envelope homeostasis in sustaining LTNS survival. All components of the Tol-Pal system were conditionally essential and deletion of *tolA* impaired the LTNS survival of *S. sonnei* at specific time points. This is consistent with previous studies demonstrating the Tol-Pal system as being important in surviving or recovering from nutrient limitation (50). The Tol-Pal system links the OM, PG and IM (50) and mediates OM invagination supporting proper envelope remodelling (51). Furthermore, Tol-Pal participates in retrograde phospholipid transport from the OM to the IM, maintaining lipid asymmetry and barrier function (52). The importance of phospholipid transport during LTNS is also highlighted by the identification of *mlaE*, encoding an IM protein of the MlaFEDB complex as a CEG in our TraDIS. Our analysis also found genes that participate in the biogenesis of OMPs (e.g., Bam complex, Sec complex, and periplasmic chaperones). We deleted *bamB,* causing defects in LTNS survival at several time points. Deletion of *bamB* in *E. coli* causes defects in envelope integrity likely through interactions with peptidoglycan synthesis (53), which may explain the observed impaired LTNS survival. Together, our results support a model in which envelope homeostasis is central for *S. sonnei* survival in starvation, likely by ensuring normal integrity/permeability that may prevent cell lysis and loss of internal components. Interestingly, components of the Tol-Pal complex were identified as important for gut colonisation in EHEC (49), and mutations in *bamB* also show reduced gut colonisation in *K. pneumoniae* (54). This again highlights the convergence of genes and pathways required for LTNS survival and enterobacterial gut colonisation, demonstrating the value of our workflow in underscoring biologically relevant pathways for *S. sonnei* pathogenesis.

We and others have observed discrepancies between results derived from TraDIS and results using deletion mutants (55–57) reinforcing the importance of validating genome-wide analysis with individual testing. For instance, despite being identified as conditionally essential by TraDIS, *metE* showed increased fitness at D14 relative to the WT. It is important to remember that TraDIS compares the fitness of each mutant within a highly mixed population and between two conditions. Our single-strain and competition assays are only partially recapitulating this complex comparison. Another important consideration is the specific mutation created. Transposon insertion can lead to partial gene disruption or polarity effects on downstream genes, which may differ from the complete loss of function derived from deletion (36,58). However, despite its limitations, TraDIS is still a valuable approach for investigating bacterial phenotypes and we found the majority of deletion mutants recapitulated the TraDIS results (35). Notably, we also found competition assays to be more sensitive to survival differences than single-strain cultures in LTNS. Bacterial fitness has generally been quantified by measuring growth in monoculture under a single stress, but competition assays are more aligned with the evolutionary biology idea of fitness as they measure comparative fitness rather than absolute fitness (59). For enteric bacteria, competition assays in a nutrient-restricted environment are an accurate reflection of their natural habitat (60).

Our study identified 418 essential genes (required under both NR and LTNS conditions) in *S. sonnei*, compared to the previously reported 498 essential genes in *S. sonnei* (61). The previous study used a different strain of *S. sonnei* (ATCC 29930) and achieved a lower number of unique insertion sites than in our current study which may contribute to the different numbers of essential genes discovered. In addition, we mapped our reads to the published sequence for strain 53G which identified 59 essential genes within an Enterobacteria phage Mu. We were unable to PCR amplify any region of this phage from our 53G isolate suggesting our isolate does not contain this phage. Therefore the number of essential genes can be reduced to 359 in our 53G strain. We anticipate the actual number of essential genes may be still lower than 359 as evidenced by the 2 pseudogenes identified as essential on pINV, a plasmid known to be non-essential for growth. Therefore, there are also likely to be small genes on the chromosome falsely identified as essential and a denser library is required to accurately identify the complement of essential genes for *S. sonnei*.

We interrogated our CEGs to determine if a single gene could explain the difference in LTNS survival between *S. sonnei* and *S. flexneri*. While *yncJ* appeared to be a promising candidate, deletion of this gene did not effect *S. sonnei* survival of LTNS. Only two unique insertion sites were found in this gene due to its small size (228 nt). Medium-density libraries such as ours can lead to overestimation of essential and conditionally essential genes (62). For this reason, a second gene which we identified as a candidate for the differential survival of *S. sonnei* and *S. flexneri, ybcO,* was not investigated further as it was a similar size (291 nt) and again contained only 2 unique insertion sites. We focussed instead on genes that had higher insertion frequencies combined with non-homology between *S. flexneri* and *S. sonnei*. The two genes we tested, *ldcA* (a PG carboxypeptidase) and *rseA* (the anti-sigma factor regulator) have roles in envelope homeostasis and were confirmed to be important for *S. sonnei* survival of LTNS. However complementation with the *S. flexneri* allele restored the survival to WT levels indicating that individually these genes were not responsible for the differential survival of *S. flexneri* and *S. sonnei* to LTNS. We suggest instead that it may be a combination of factors that results in the differential survival in LTNS of these two *Shigella* species rather than a single gene difference.

Taken together, here we have demonstrated significant differences in the abilities of *S. flexneri* and *S. sonnei* to survive nutrient starvation. This finding provides new evidence underscoring the divergence between these two *Shigella* species that may contribute to explaining their different epidemiology and pathogenesis.

## Materials and methods

### Bacterial strains and growth conditions

Bacterial strains used in this study are listed in Table 3. *Shigella* strains were routinely grown in Tryptic Soy Broth (TSB) or Tryptic Soy Agar (TSA) plates supplemented with 0.01% Congo Red, unless otherwise stated. Congo Red allows identification of colonies that harbour the large virulence plasmid (LVP) (63,64). *E. coli* strains were grown in Lysogeny Broth (LB) or Lysogeny Agar (LA) plates. Cultures were incubated at 37°C overnight. If required, antibiotics were added to the media with the following concentrations: chloramphenicol (Cm, 30 µg/ml), gentamicin (Gm, 20 µg/ml), kanamycin (Kn, 50 μg/ml).

**Table 3.**
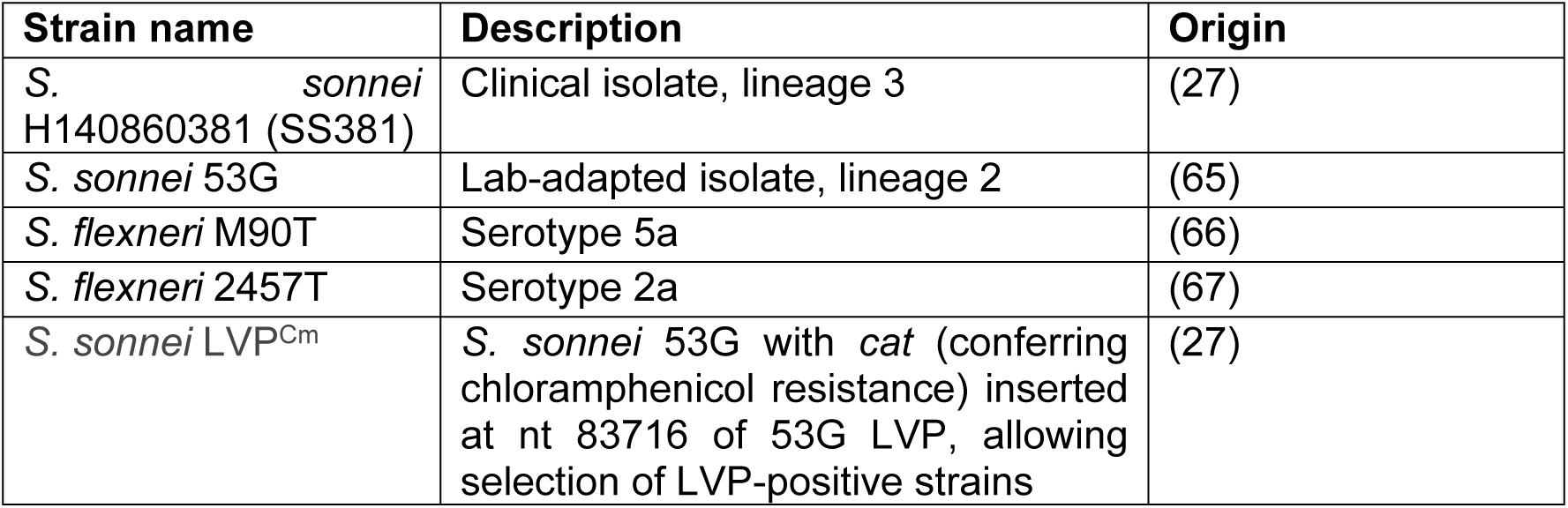
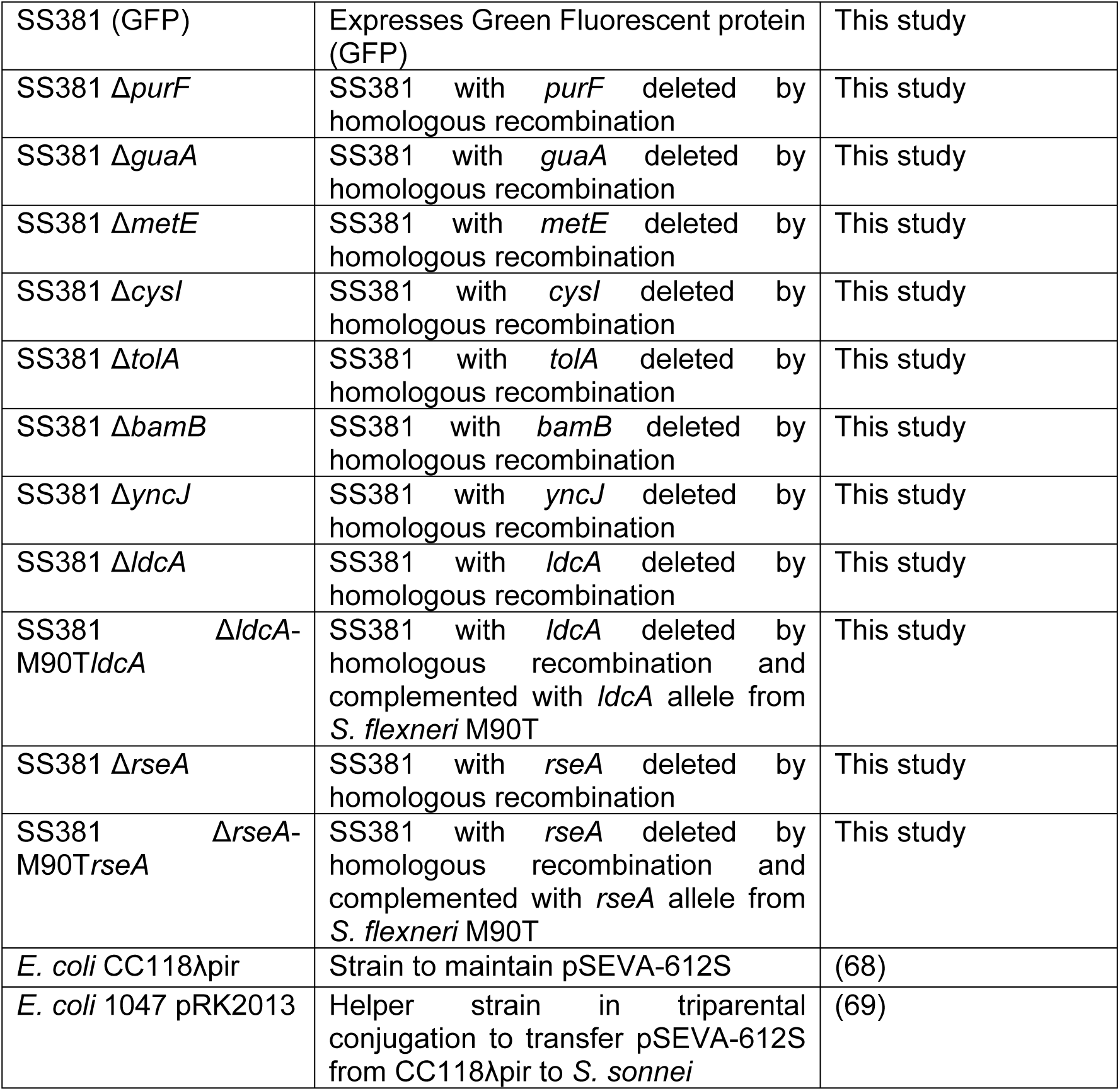
Bacterial strains table.

### Long-term nutrient starvation (LTNS) cultures

To compare the survival of *S. sonnei* and *S. flexneri* strains in long-term nutrient starvation (LTNS) conditions we used minimal media M9 (33.7 mM Na₂HPO₄, 22 mM KH₂PO₄, 8.55 mM NaCl, 9.35 mM NH₄Cl) supplemented with 2 mM MgSO₄ and 0.1 mM CaCl. Strains were first grown in TSB to the exponential phase and washed with PBS prior to inoculation in M9. Independent M9 cultures of each strain at a concentration of 10^6^ cells/ml were incubated at 23 ± 2°C, and the number of CFUs was monitored over a period of 18 days. At specific timepoints, aliquots were collected, and serial dilutions were plated onto TSA to quantify for viable CFUs.

To test for differences in the survival in LTNS between different strains derived from *S. sonnei* SS381, we additionally carried out competition experiments. Strains were first grown independently in TSB to the exponential phase, washed with PBS, and inoculated together in M9 at a concentration of 10^6^ cells/ml per strain. Cultures were incubated at 23 ± 2°C, aliquots were taken over 14 days at different time points, and CFUs were quantified. To allow rapid strain differentiation in competition assays, these were carried out using pairs of strains in which one was GFP-labelled. We used the blue-light transilluminator Safe Imager 2.0 (Invitrogen) for visual discrimination.

### Generation of transposon mutant libraries and sequencing

We constructed a transposon mutant library in *S. sonnei* strain 53G:LVP^Cm^ using transposomes prepared from mini-Tn5 and EZ-Tn5 Transposase (Epicentre). Transformants were collected from TSA plates containing 0.01% Congo Red, 8.5 μg/ml Cm, and 50 μg/ml Kn. The bacteria were resuspended at OD_600_ of 100 and immediately stored in 1ml aliquots at -80°C. The number of bacteria stored was retrospectively estimated by CFU determination.

Individual aliquots of the transposon library (10^9^ cfu/ml) were thawed and added to M9 and subjected to LTNS. After 18 days DNA was extracted from each LTNS culture and from a frozen aliquot of the transposon library via phenol-chloroform extraction.

Two micrograms of DNA from each gDNA preparation were used to prepare TraDIS transposon-specific sequencing libraries following the protocol described in the TraDIS Toolkit method (70), with TraDIS adapter and primers as described previously (70). The resulting DNA was sequenced on an Illumina MiSeq platform using a MiSeq reagent kit V2 (50 cycles) (Illumina, USA).

The analysis of TraDIS sequencing results was carried out using the Bio-TraDIS pipeline (https://github.com/sanger-pathogens/Bio-Tradis) as described previously (70). Processed reads were mapped to the reference genome (chromosome HE616528, plasmid A HE616529, plasmid B HE616530, plasmid C HE616531 and plasmid E HE616532). The input pool achieved read counts of 2,430,678 and 281800 for the chromosome and large virulence plasmid respectively. Statistical analysis was carried out using R v.3.2.3 included in the Bio-Tradis pipeline. The *bacteria_tradis* command was run using default settings, except the following parameters: --smalt_r 0 -m 0 and -t TAAGAGACAG. These settings enable mapping of multisite mapping reads, which would otherwise be filtered out, and ensure mapping of reads containing the expected transposon tag. The resulting insertion count plots (insert_site_plot.gz files, one per each replicon, i.e., NC_016822, NC_016823, NC_016824, NC_016833, and NC_016834) were then processed with the *tradis_gene_insert_sites* command against the relevant annotated reference replicon. *-trim3* was set to *0.1* to trim reads at the 3′ end, as many essential genes tolerate insertions towards the end of the coding sequence (71). A summary of genes that tolerated (i.e., non-essential genes) and did not tolerate (i.e., essential genes) insertions was produced by running the *tradis_essentiality.R* command on the resulting tradis_gene_insert_sites.csv files. Consensus essential genes were defined as genes where no insertions were identified in any of the samples. For the analysis of conditionally essential genes in long-term nutrient starvation, the logFC of read counts and the False Discovery Rate (FDR)-adjusted *p*-value (*q*-value) for each gene were evaluated by using the *tradis_comparisons.R* script. Genes with significantly enriched transposon insertions were selected by cutoff as logFC ≥2 and *q*-value ≤0.05.

### GO categories

PANTHER (Protein ANalysis THrough Evolutionary Relationships) Classification System (https://pantherdb.org/) was used to classify genes that were identified by TraDIS according to their function using GO terms (GO Phylogenetic Annotation Project). Genes were attributed to significant functional categories (p-value < 0.05) using Fisher’s test and FDP correction.

### Mutagenesis and complementation

The plasmid vector pSEVA-612S served as a backbone for both mutagenesis and complementation constructs for the genetic modifications of the *S. sonnei* strain SS381.

The insertion of the green fluorescence protein (GFP) was made immediately downstream of *glmS*, previously shown to be a neutral site (38). For this purpose, the GFP-encoding gene was amplified from pULTRA-GFP and cloned into pSEVA-612S, flanked by ±500 bp upstream and downstream regions surrounding the insertion site downstream of *glmS*. Similarly, for the construction of deletion mutants, ±500 bp upstream and downstream regions flanking the target gene were cloned into plasmid pSEVA-612S. For the complementation of Δ*ldcA* and Δ*rseA* with *S. flexneri* alleles, *ldcA* and *rseA* were amplified from *S. flexneri* M90T and cloned into pSEVA-612S flanked by corresponding ±500 bp upstream and downstream regions of SS381. All PCRs were carried out using Q5 DNA polymerase (New England Biolabs, NEB) to ensure high fidelity amplification and cloning constructs were obtained by Gibson Assembly (NEB) or by restriction-ligation with T4 DNA ligase (NEB).

The resulting mutagenesis/complementation plasmids were then introduced into *E. coli* CC118λpir and then mobilised into *S. sonnei* by triparental conjugation with helper strain *E. coli* 1047 pRK2013. To resolve *S. sonnei* merodiploids, the I-SceI endonuclease encoded on plasmid pACBSR was induced by adding 0.4% (w/v) L-arabinose to the culture. After homologous recombination occurred, mutagenesis/complementation was confirmed by PCR screening and Sanger sequencing (Eurofins Genomics). Primers used in this study are listed in Supplementary Table 5.

### Comparative sequence analysis

To identify *S. sonnei* specific variants potentially contributing to the differing survival between *S. sonnei* and *S. flexneri* strains, we compared the amino acid sequences of all conditionally essential genes identified by TraDIS in *S. sonnei* 53G with their counterparts in *S. flexneri* serotype 5a strain M90T and *S. flexneri* serotype strain 2a 2457T. Pairwise alignments were performed using BLASTp (https://blast.ncbi.nlm.nih.gov/Blast.cgi) with default parameters. The *S. sonnei* 53G protein sequences were used as queries, and the following reference genomes were used as subjects: *S. flexneri* serotype 5a strain M90T (NCBI RefSeq: NZ_CP037923.1), *S. flexneri* 2a strain 2457T (GenBank: AE014073.1), and *S. sonnei* strain 381 (GenBank: GCA_001248585.1). Genes were considered of interest if they were absent or non-homologous in both *S. flexneri* genomes and conserved (100% identical) between *S. sonnei* 53G and *S. sonnei* 381.

### Statistical analysis

Statistical analyses were conducted using GraphPad Prism (v 10.4.1). The specific tests applied to each dataset are detailed in the corresponding figure legends. Statistical significance is denoted as follows: ns (not significant), * (P ≤ 0.05), ** (P ≤ 0.01), *** (P ≤ 0.001), and **** (P ≤ 0.0001).

## Supplementary materials

### Bacterial growth kinetics

Overnight cultures of *S. sonnei* strains in TSB were diluted in fresh media to an OD_600_ of ∼0.1. 100 μL of each suspension were added to a 96-well plate in triplicate. The plate was incubated at 37°C with the OD_600_ measured every 10 minutes over 12 hours with shaking (200rpm).

### Protein sequence alignment

Multiple sequence alignments of amino acid sequences of interest were performed using Clustal Omega (https://www.ebi.ac.uk/jdispatcher/msa/clustalo) with default parameters.

## Acknowledgments

We would like to thank the Wellcome Trust Sanger Institute sequencing facility for TraDIS library preparation and sequencing. This work was supported by a Medical Research Council grant MR/X00080X/1 and a Royal Society grant RG130136 (A.C). P.B.L was supported by a Wellcome Trust PhD studentship (220057/Z/19/Z). V.T is currently supported by a Medical Research Council New Investigator Research Grant (MR/Z504178/1).

---

**Supplementary Fig 1. Non-competitive and competitive LTNS survival of WT SS381 in comparison to SS381 (GFP)**

A) Non-competitive LTNS survival of WT SS381 and SS381 (GFP). Data show mean values ± SEM of n = 3 biological replicates. Log_10_-transformed data was analysed by 2-way ANOVA with main column effect compared by Tukey’s multiple comparison. ns = non-significant (p > 0.05). B) Competition assays between WT SS381 and SS381 (GFP). Data show mean values ± SEM of n=3 biological replicates. A two-way ANOVA with Šídák’s multiple comparisons was used to compare mean values at different time points. ns = non-significant (p > 0.05).

**Supplementary Fig 2. Bacterial growth in nutrient rich (NR) conditions of strains used in this study**

Growth was measured by recording optical density values at 600 nm (OD_600_) every 10 minutes for a period of 12 hours in TSB at 37°C. Data show mean values of 2 biological repeats.

**Supplementary Fig 3. Alignment of LdcA and RseA protein sequences from *S. sonnei* and *S. flexneri***

A) Amino acid alignment of LdcA from *S. sonnei* 53G, *S. sonnei* 381, *S. flexneri* M90T, and *S. flexneri* 2457T showing three non-conservative amino acid differences (highlighted in blue). B) Amino acid alignment of LdcA from *S. sonnei* 53G, *S. sonnei* 381, *S. flexneri* M90T, and *S. flexneri* 2457T showing one non-conservative (highlighted in blue) and located within the N-terminal σ^E-binding domain. Alignments were performed using Clustal Omega.

